# m^6^A-binding YTHDF proteins promote stress granule formation by modulating phase separation of stress granule proteins

**DOI:** 10.1101/694455

**Authors:** Ye Fu, Xiaowei Zhuang

## Abstract

Diverse RNAs and RNA-binding proteins form phase-separated, membraneless granules in cells under stress conditions. However, the role of the prevalent mRNA methylation, m^6^A, and its binding proteins in stress granule (SG) assembly remain unclear. Here, we show that m^6^A-modified mRNAs are enriched in SGs, and that m^6^A-binding YTHDF proteins are critical for SG formation. Depletion of YTHDF1/3 inhibits SG formation and recruitment of m^6^A-modified mRNAs to SGs. Both the N-terminal intrinsically disordered region and the C-terminal m^6^A-binding YTH domain of YTHDF proteins are crucial for SG formation. Super-resolution imaging further reveals that YTHDF proteins are in a super-saturated state, forming clusters that reside in the periphery of and at the junctions between SG core clusters, and promote SG phase separation by reducing the activation energy barrier and critical size for condensate formation. Our results reveal a new function and mechanistic insights of the m^6^A-binding YTHDF proteins in regulating phase separation.

## Main

RNA-protein (RNP) granules are phase-separated, membraneless granules that play important roles in epigenetic and post-transcriptional regulations^1–6^. Stress granules (SGs) are RNP granules that assemble under various cellular stress conditions, such as oxidative, osmotic, or heat-shock stress, and regulate messenger RNA (mRNA) translation and degradation^1,4,5^. Defects in SG dynamics are associated with various diseases such as neurodegenerative disorders, cancers, viral infections, and autoimmune diseases^7,8^.

RNAs and RNA-interacting proteins are crucial components of SGs^9–12^. *N*^6^-methyladenosine (m^6^A) is the most abundant internal mRNA modification^13–16^, and three of the major m^6^A-binding proteins, YTHDF1-3 (ref. ^15^), are implicated in the SG proteome and can interact with SG components^12,17–20^. However, some key components of SGs, including the SG core proteins, G3BP1/2 (ref. ^21^), and their binding partners^22^, preferentially bind to unmodified RNA instead of m^6^A-modified RNAs in specific sequence contexts^23^. These opposing binding behaviors of SG proteins to m^6^A-modified RNAs and m^6^A-binding proteins raise the important question of whether m^6^A-modified RNAs and m^6^A-binding proteins play a role in SG formation.

Here, we studied the localization of m^6^A-modified mRNAs and m^6^A-binding YTHDF proteins in mammalian cells, and identified the m^6^A-binding YTHDF proteins as key regulators for SG formation. We observed an enrichment of m^6^A-modified mRNAs in SGs. Depletion of YTHDF1/3 proteins substantially inhibited SG formation and prevented enrichment of mRNA m^6^A signal in SGs. Both the N-terminal intrinsically-disordered region (IDR) and C-terminal m^6^A-binding YTH domain of YTHDF proteins were crucial for SG formation. Super-resolution imaging further revealed that YTHDF1 was in a super-saturated state, forming clusters that connect SG core clusters, and promoted phase separation of the SG core protein G3BP1 by lowering the activation energy barrier and reducing the critical size for G3BP1 condensate formation.

## Results

### Imaging m^6^A-modified mRNA in mammalian cells

To examine the subcellular distribution of m^6^A-modified mRNAs in mammalian (U-2 OS) cells, we developed an immunofluorescence protocol that specifically labels m^6^A-modified mRNAs (Supplementary Fig. 1a, b) and validated the labeling specificity for m^6^A using gene-edited U-2 OS cells in which a key m^6^A methyltransferase component METTL3 is knocked out (Supplementary Fig. 1c). It has been shown previously that the amount of m^6^A in mRNAs is reduced by 50 - 60% in METTL3 knockout cells^24^ and indeed, our immunofluorescence signal for mRNA m^6^A was reduced by ~60% in these knockout cells (Supplementary Fig. 1c).

Using this imaging approach, we observed several notable features for the distribution of mRNA m^6^A in unstressed cells: First, the normalized intensity of the mRNA m^6^A signal (normalized by polyA signal) was substantially higher in the cytoplasm than in the nucleus (Supplementary Fig. 1d), which is consistent with the effect of m^6^A on promoting nuclear export of mRNAs^25^. Second, in the cytoplasm, in addition to the diffusively distributed signals, we observed an enrichment of m^6^A in processing bodies (P-bodies) (Supplementary Fig. 1e).

### Enrichment of m^6^A-modified mRNA in SGs

To study the localization of mRNA m^6^A under stressed conditions, we imaged mRNA m^6^A and polyA signals simultaneously in U-2 OS cells under oxidative stress induced by NaAsO_2_ treatment. NaAsO_2_ treatment induced the formation of numerous SGs in the cytoplasm, marked by an SG core protein G3BP1 (Fig. 1a)^22^. We observed strong signals of both mRNA m^6^A and polyA in SGs (Fig. 1a). Quantitatively, polyA showed ~3-fold enrichment in SGs as compared to elsewhere in the cytoplasm whereas m^6^A showed more than 4-fold enrichment in SGs (Fig. 1a), suggesting that m^6^A-modified mRNAs have a higher tendency to associate with SGs than unmethylated mRNAs. We also observed strong m^6^A signals but not polyA signals in P-bodies (Fig. 1a), which is consistent with previous observations that the m^6^A-binding protein YTHDF2 localizes in P-bodies^26^ and that deadenylation of mRNAs is a prerequisite for P-body formation^27^.

**Fig. 1.**
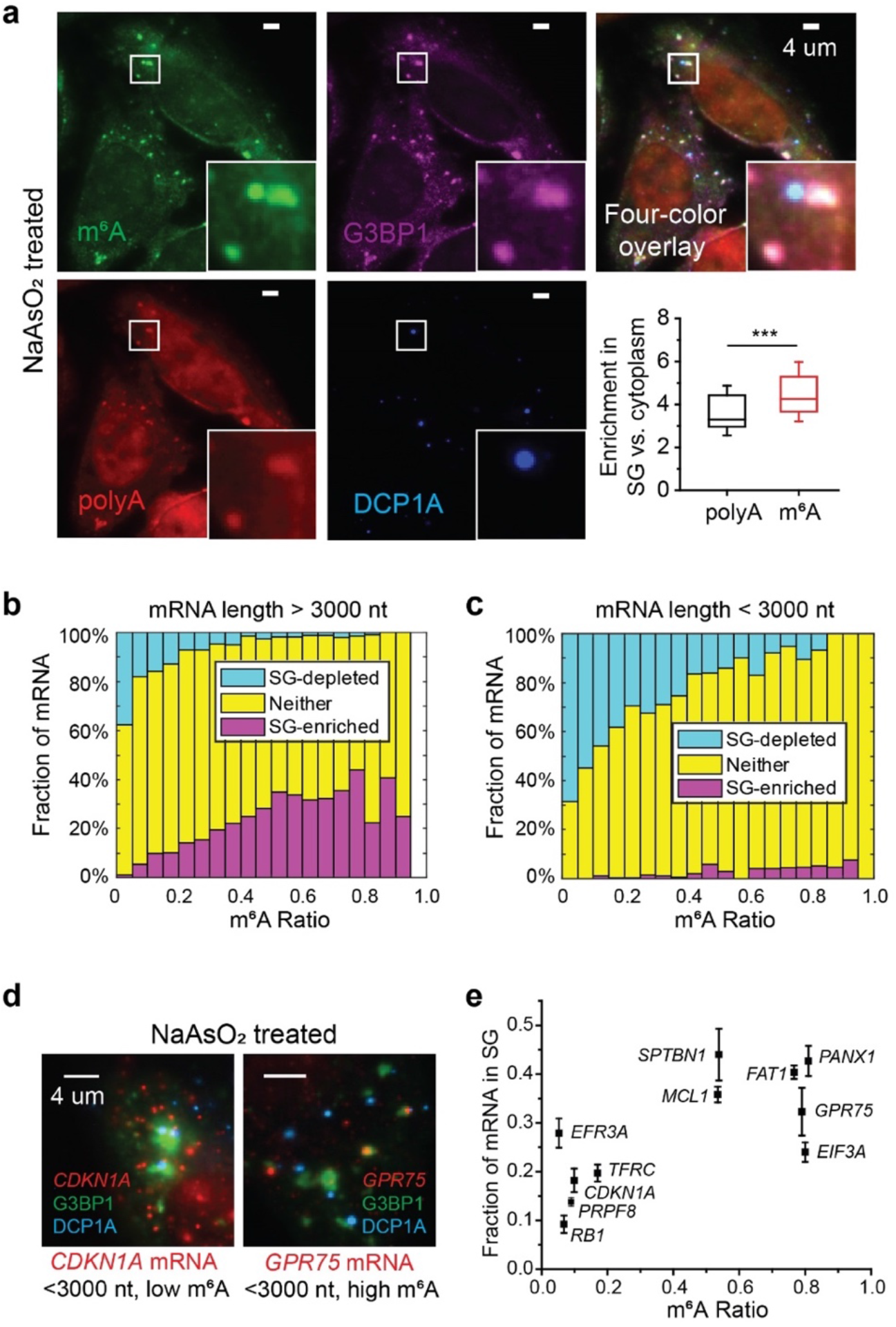
m^6^A-modified mRNAs are enriched in stress granules (SGs) in U-2 OS cells under oxidative stress. **a**, Immunofluorescence staining of mRNA m^6^A in U-2 OS cells treated with 0.5 mM NaAsO_2_ for 30 min shows enrichment of m^6^A signal in SGs and P-bodies. SG marker G3BP1 and P-body marker DCP1A are detected using immunofluorescence and polyA signal is detected using FISH. The bottom right panel shows box plots of the enrichment of m^6^A signal and polyA signal in SGs. The box plots show the median (middle lines), 25%-75% quartiles (boxes), and standard deviation (error bars). n =55 cells, from 3 independent experiments. *** P < 0.001 using unpaired Mann–Whitney U test. **b**, **c**, Transcriptome-wide analysis shows the fractions of mRNAs that are enriched in SGs (magenta), depleted in SGs (cyan), and neither enriched nor depleted in SGs (yellow), as a function of the m^6^A ratio for individual mRNAs that are longer than 3000 nt (**b**), or shorter than 3000 nt (**c**). The m^6^A ratio is defined as the percentage of transcripts that contain m^6^A. n = 9049 genes in total. **d**, Examples of smFISH images showing that the mRNA with a higher m^6^A ratio tends to have a higher degree of enrichment in SGs. Cells treated with 0.5 mM NaAsO_2_ for 30 min to induce oxidative stress. **e**, The fraction of mRNA localized in SG measured using smFISH versus the m^6^A ratio for individual mRNAs. Error bars represent mean ± SEM. n = 32 cells (RB1), 60 cells (PRPF8), 50 cells (CDKN1A), 70 cells (TFRC), 24 cells (EIF3A), 32 cells (EFR3A), 13 cells (SPTBN1), 87 cells (MCL1), 159 cells (FAT1), 43 cells (PANX1), 51 cells (GPR75), from 3 independent experiments each.

We then analyzed the relationship between the SG-enrichment of mRNAs determined by SG RNA sequencing^28^ and the m^6^A methylation ratio of mRNAs (defined as the fraction of transcripts that harbor m^6^A)^29^ for individual genes. We observed a strong positive correlation between m^6^A ratio and SG-enrichment and negative correlation between m^6^A ratio and SG-depletion for relatively long (>3000 nt) mRNAs (Fig. 1b). For relatively short (<3000 nt) mRNAs, few of them showed enrichment in SGs but the strong negative correlation between m^6^A ratio and SG-depletion maintained (Fig. 1c). We further validated the localization of various mRNAs with different m^6^A ratios using single-molecule fluorescent *in*-*situ* hybridization (smFISH)^30,31^, which also showed a higher tendency of SG-enrichment for mRNAs that are more heavily modified with m^6^A (Fig. 1d, e). We observed a similar trend for mRNA enrichment in other types of RNP granules, including heat-shock induced SGs^32^, ER-stress induced SGs^32^, and P-bodies^33^ (Supplementary Fig. 2). Taken together, our results indicate that a broad range of m^6^A-modified mRNAs are enriched in SGs.

### RNA m^6^A-binding YTHDF proteins are critical for SG formation

To understand the mechanism underlying the association between m^6^A-modified mRNAs and SGs, we examined the roles of m^6^A-binding YTHDF proteins in SG assembly. We observed strong colocalization of endogenous YTHDF1/3 proteins with SGs, but not with P-bodies, which are frequently found adjacent to SGs (Fig. 2a, c; Supplementary Fig. 3a, b, c). YTHDF2, instread, showed colocalization with both SGs and P-bodies (Fig. 2b; Supplementary Fig. 3d). Notably, knockdown of either YTHDF1 or YTHDF3, but not YTHDF2, substantially reduced the number of SGs in NaAsO_2_-treated cells (Fig. 3a, b). Double knockdown of YTHDF1 and YTHDF3 largely abolished the formation of SGs (Fig. 3a, b). The reduction in SG formation upon YTHDF1/3 knockdown was accompanied by a substantial reduction of both polyA and m^6^A signals in SGs (Fig. 3c).

**Fig. 2.**
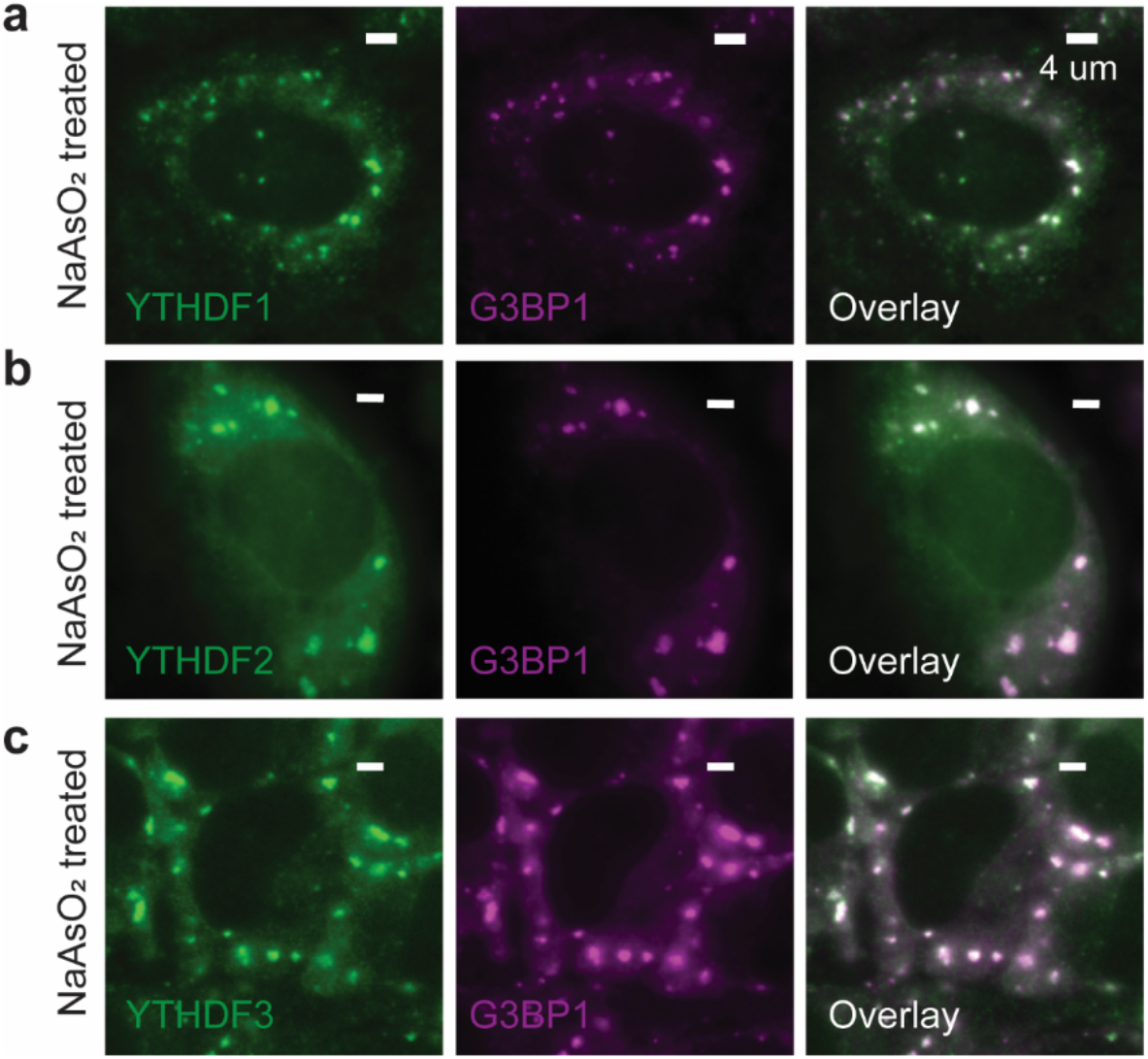
YTHDF proteins are enriched in SGs. Conventional immunofluorescence images of endogenous YTHDF1-3, and G3BP1 show that YTHDF1 (a), YTHDF2 (b) and YTHDF3 (c) are enriched in SGs after oxidative stress.

**Fig. 3.**
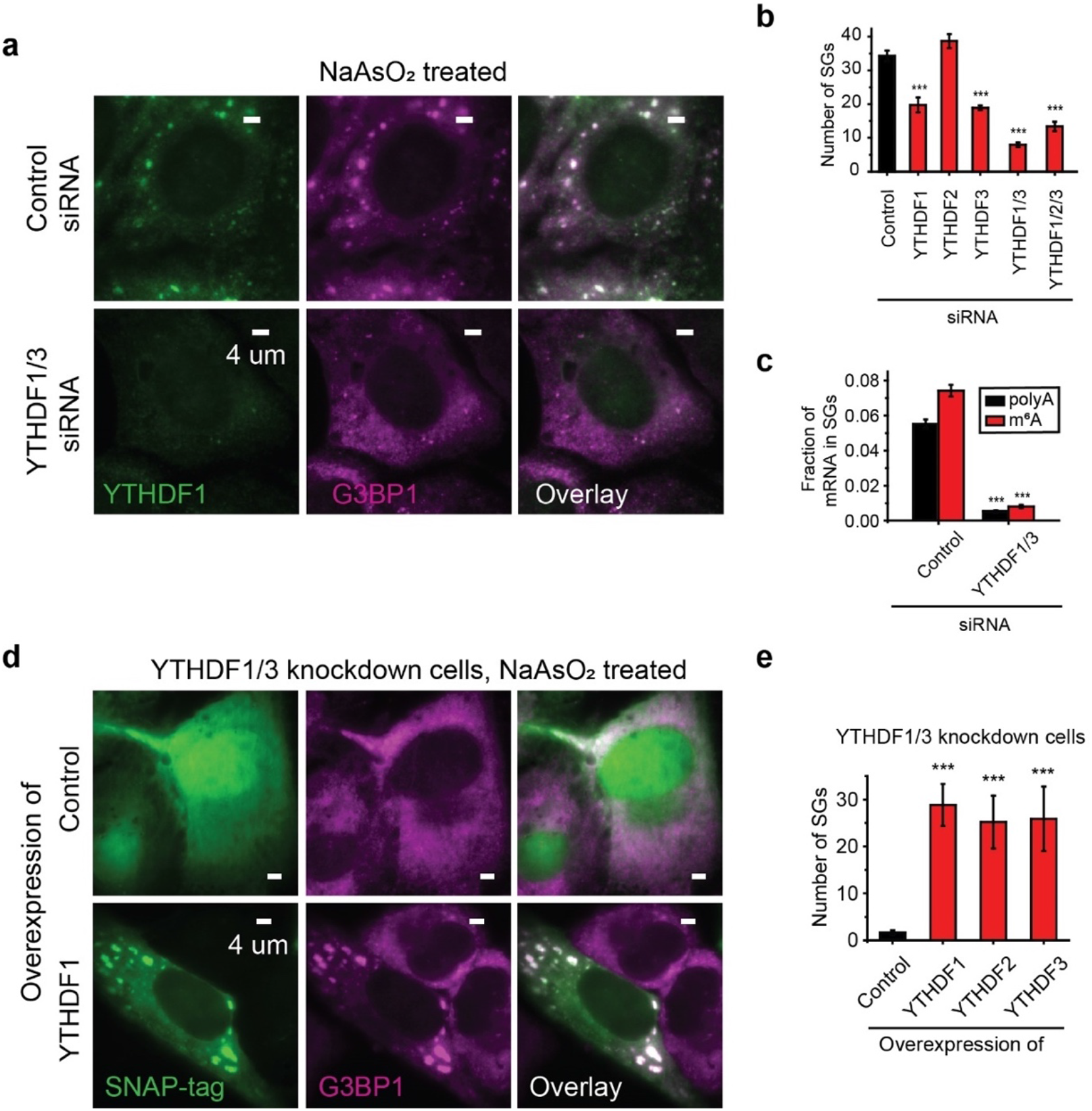
YTHDF proteins promote SG formation. **a**, Two-color immunofluorescence images of YTHDF1 and G3BP1 show the disappearance of large SGs upon YTHDF1 and YTHDF3 double siRNA knockdown. The upper panels show the images for cells treated with control (scrambled) siRNA and the lower panels show the images of the YTHDF1/3 double knockdown cells. **b**, Quantification of the number of SGs per cell for U-2 OS cells treated by control siRNA, single knockdown cells treated with YTHDF1, YTHDF2 or YTHDF3 siRNA, YTHDF1/3 double knockdown cells, and YTHDF1/2/3 triple knockdown cells. Oxidative stress in these cells was induced by 0.5 mM NaAsO_2_ treatment for 40 min. Bar graph represents mean ± SEM here. *** P < 0.001 compared with control; Unpaired Mann–Whitney U test. n = 256 cells (control siRNA), n = 107 cells (YTHDF1 siRNA), n = 221 cells (YTHDF2 siRNA), n = 243 cells (YTHDF3 siRNA), n = 203 cells (YTHDF1/3 siRNA), n = 42 (YTHDF1/2/3 siRNA), from 3 independent experiments each. **c**, Fraction of polyA (black) and m^6^A (red) signals in SGs in cells treated with control siRNA as well as in YTHDF1/3 double knockdown cells. Bar graph represents mean ± SEM here. *** P < 0.001 compared with control; Unpaired Mann–Whitney U test. n = 256 cells (control siRNA), n = 203 cells (YTHDF1/3 siRNA), from 3 independent experiments each. **d**,**e**, Overexpression of full-length YTHDF1, YTHDF2, and YTHDF3 proteins rescues the SG formation in YTHDF1/3 knockdown cells. All constructs are tagged with SNAP at the C-terminal end. Overexpressed proteins were imaged using a fluorescent dye that labels the SNAP-tag. **d**, Two color images of SNAP-tag, detected by dye molecules conjugated to SNAP, and G3BP1, detected by immunofluorescence, for cells expressing a control SNAP-tag plasmid that does not contain YTHDF (upper panels) and for cells expressing SNAP-YTHDF1 (lower panels). Cells were treated by 0.5 mM NaAsO_2_ for 25 min to induce oxidative stress. **e**, Quantification of the number of SGs per cell for YTHDF1/3 double knockdown cells overexpressing the control SNAP-tag and for YTHDF1/3 double knockdown cells overexpressing SNAP-YTHDF1, SNAP-YTHDF2 or SNAP-YTHDF3. Bar graph represents mean ± SEM here. *** P < 0.001 compared with control; Unpaired Mann–Whitney U test. Bar graph represents mean ± SEM. n = 36 cells (Control), n = 33 cells (YTHDF1), n = 25 cells (YTHDF2), n = 19 cells (YTHDF3), from 3 independent experiments each.

To further confirm the effect of YTHDF proteins in SG formation, we overexpressed individual YTHDF proteins to compensate for the effect of YTHDF1/3 knockdown. Overexpression of YTHDF1/3 indeed restored SG formation (Fig. 3d, e). Interestingly, although knocking down endogenous YTHDF2 did not show a substantial effect on SG formation (Fig. 3b), overexpression of this protein also restored SG formation in YTHDF1/3 knockdown cells, potentially because the effect of YTHDF2 is weak at the physiological concentration but substantial at elevated concentrations.

### YTHDFs’ effect in SG formation requires both N-IDR and YTH domain

Many RNA-binding proteins in SGs possess intrinsically disordered regions (IDRs) and/or prion-like domains (PLDs), which can promote liquid-liquid phase separation^34^. We analyzed the amino acid sequences and secondary structures of YTHDF proteins (Fig. 4a; Supplementary Figs. 4, 5) using three algorithms (NetSurfP-2.0 for secondary structure prediction^35^, PLAAC (Prion-Like Amino Acid Composition) for PLD detection^36^, and PONDR-VSL2, Predictor of Natural Disordered Regions based on Various training data for Short and Long disordered sequences^37^). Secondary structure and disordered region predictions showed that while the C-terminal RNA-binding YTH domain contains defined structures, the remaining parts of YTHDF proteins are largely disordered (Fig. 4a, Supplementary Fig. 5). PLD analysis further identified a sub-region in the disordered region that has consistently high PLD scores and is Pro(P)/Gln(Q)-rich (Fig. 4a, Supplementary Fig. 5) – we referred to this region as the P/Q-PLD and the remaining (N-terminal) part of the disordered region as the N-IDR (Fig. 4a, Supplementary Fig. 5).

**Fig. 4.**
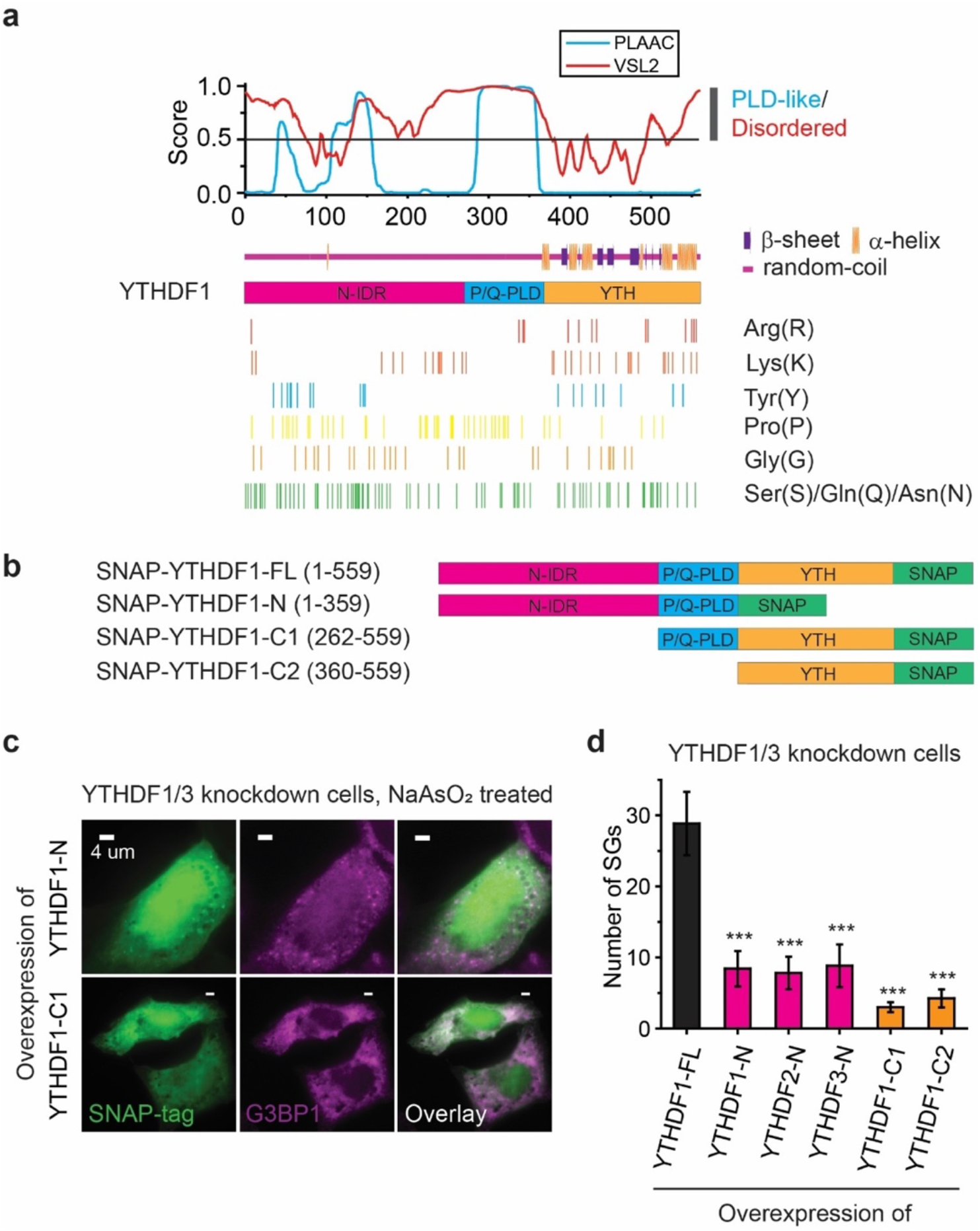
Both the N-terminal intrinsically disordered region (N-IDR) and the C-terminal YTH domain are essential for YTHDF’s role in promoting SG formation. **a**, Amino acid composition and predictions of IDRs, prion-like domains (PLDs), and secondary structures in YTHDF1 protein. The likelihood scores for being disordered predicted by PONDR-VSL2 (red) and being PLD-like predicted by PLAAC (cyan) for each amino acid are plotted in the upper panel. The secondary structure prediction by NetSurfP-2.0 is shown immediately below the disordered and PLD-like likelihood scores. Locations for several conserved amino acids in all three YTHDF proteins that are important for IDR/PLD properties are marked. **b**, Different truncated YTHDF1 constructs tagged with SNAP at the C-terminal end. **c**,**d**, Overexpression of different YTHDF protein fragments in YTHDF1/3 knockdown cells shows that truncated YTHDF constructs lacking the N-IDR or YTH domain cannot rescue SG-formation. **c**, Two color images of SNAP-tag, detected by dye molecules conjugated to SNAP, and G3BP1, detected by immunofluorescence, for cells expressing a YTHDF1 fragment lacking the YTH domain (upper panels) or the N-IDR (lower panels). The cells are treated with 0.5 mM NaAsO_2_ for 25 min to induce oxidative stress. **d**, Quantification of the number of SGs per cell for YTHDF1/3 double knockdown cells overexpressing the full-length YTHDF1 (black), or YTHDF1/3 double knockdown cells overexpressing YTHDF1/2/3 fragments lacking YTH (magenta), YTHDF1 fragment lacking N-IDR, or YTHDF1 fragment lacking both N-IDR and P/Q-PLD (orange). Bar graph represents mean ± SEM here. *** P < 0.001 compared with full-length YTHDF1; Unpaired Mann–Whitney U test. n = 33 cells (YTHDF1-FL), n = 19 cells (YTHDF1-N), n = 23 cells (YTHDF2-N), n = 12 cells (YTHDF3-N), n = 23 (YTHDF1-C1), n = 16 cells (YTHDF1-C2), from 3 independent experiments each.

Notably, whereas overexpressing full-length YTHDF proteins rescued SG formation in YTHDF1/3 knockdown cells (Fig. 3d, e), overexpressing YTHDF fragments that miss either the N-IDR or the YTH domain did not rescue SG formation in YTHDF1/3 knockdown cells (Fig. 4b, c, d), indicating that both domains are essential for the YTHDF’s role in promoting SG formation. Surprisingly, although some PLDs could promote protein aggregation^34^, only overexpressing a fragment that contained both the P/Q-PLD and the YTH domain also did not rescue SG formation in YTHDF1/3 knockdown cells (Fig. 3d).

### Blocking m^6^A-binding of YTHDF impedes SG formation

To test whether the interactions between YTHDF proteins and the m^6^A modification in RNAs are important for SG formation, we constructed a dominant-negative mutant of YTHDF1 that harbors a D401N mutation in the YTH domain (Fig. 5a). This mutation is known to increase the m^6^A binding affinity of YTH domain by 10-fold^38^ and therefore we expect overexpression of this mutant in cells to inhibit the m^6^A binding of the endogenous YTHDF proteins. We further replaced the N-IDR in the mutant by a Cry2Olig domain that can undergo blue-light-induced oligomerization^39^. Overexpression of this dominant-negative mutant in U-2 OS cells partially disrupted the formation of SGs in NaAsO_2_-treated cells, but overexpression of the control construct harboring only the Cry2Olig region did not affect SG formation (Fig. 5b, c). These results suggest that the interactions between YTHDFs and m^6^A facilitate SG formation.

**Fig. 5.**
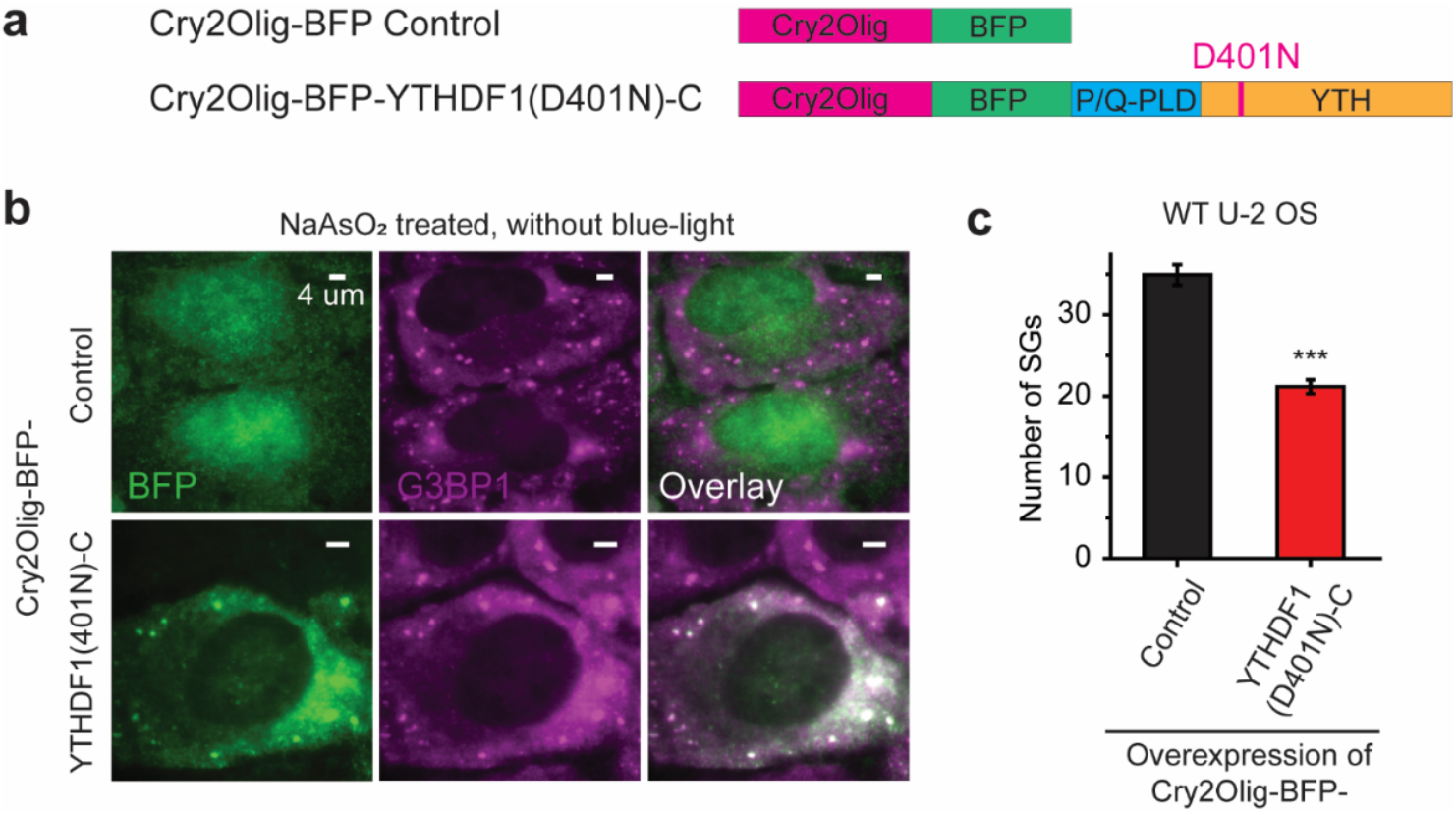
Inhibiting m^6^A-binding of YTHDF proteins impairs SG formation. **a**, A construct that contains Cry2Olig, a blue-light inducible oligomerization domain, and BFP (upper), and a construct that contains Cry2Olig, BFP, and a mutated YTHDF1 fragment (YTHDF1(D401N)-C), which harbors a single amino acid mutation with 10-fold higher binding affinity to m^6^A than wild-type YTH domains and lacks the N-IDR (lower). **b**, **c**, Formation of SGs is reduced in cells overexpressing construct containing the YTHDF1(D401N)-C mutant. **b**, Two color images of BFP and G3BP1 for cells expressing a control plasmid that contains Cry2Olig-BFP but does not contain the YTHDF1(D401N)-C mutant (upper panels) and for cells expressing Cry2Olig-BFP-YTHDF1(D401N)-C. Cells are treared with 0.5 mM NaAsO_2_ for 25 min to induce oxidative stress. **c**, Quantification of the number of SGs per cell for U-2 OS cells overexpressing Cry2Olig-BFP and U-2 OS cells overexpressing Cry2Olig-BFP-YTHDF1(D401N)-C. The cells were treated with 0.5 mM NaAsO_2_ for 25 min. Bar graph represents mean ± SEM. n = 167 cells (Cry2Olig-BFP), n = 123 cells (Cry2Olig-BFP-YTHDF1(D401N)-C), from 3 independent experiments each. ***: p < 0.001 by unpaired Mann-Whitney U test.

We then induced the oligomerization of the Cry2Olig-YTHDF1(D401N)-C mutant using blue light in unstressed cells. Although blue light illumination caused clustering of this mutant protein, which colocalized with P-bodies marked by DCP1A, this clustering of Cry2Olig-YTHDF1(D401N)-C did not induce SG formation (Supplementary Fig. 6), presumably because self-interaction of the N-terminal region of the m^6^A-bound YTHDF proteins is not sufficient for SG formation. This result is consistent with our observation that expression of the construct containing the P/Q-PLD and YTH domains, but missing the N-IDR, did not enhance SG formation in YTHDF1/3 knockdown cells. As will be discussed below, our results suggest that the intermolecular interactions between the N-IDR and YTH domains of YTHDF proteins are important for promoting SG formation.

### YTHDF proteins reduce the critical radius and activation energy for condensate formation of SG core protein G3BP1

To further understand how YTHDF proteins promote SG formation, we performed super-resolution STORM imaging^40^ of the endogenous G3BP1 protein in U-2 OS cells in the presence and absence of YTHDF1/3. The high-resolution of our images (~20 nm resolution) revealed that G3BP1 formed small clusters with size up to 200 nm in unstressed cells (Fig. 6a, b). Because of their small sizes and high density in cells, these G3BP1 clusters were not visible using diffraction-limited imaging. Upon addition of NaAsO_2_ to induce oxidative stress, the sizes of G3BP1 clusters increased substantially, with some clusters reaching 600 nm in size (Fig. 6a, b). Knocking down of YTHDF1/3 substantially reduced the sizes of the G3BP1 clusters in NaAsO_2_-treated cells (Fig. 6a, b).

**Fig. 6.**
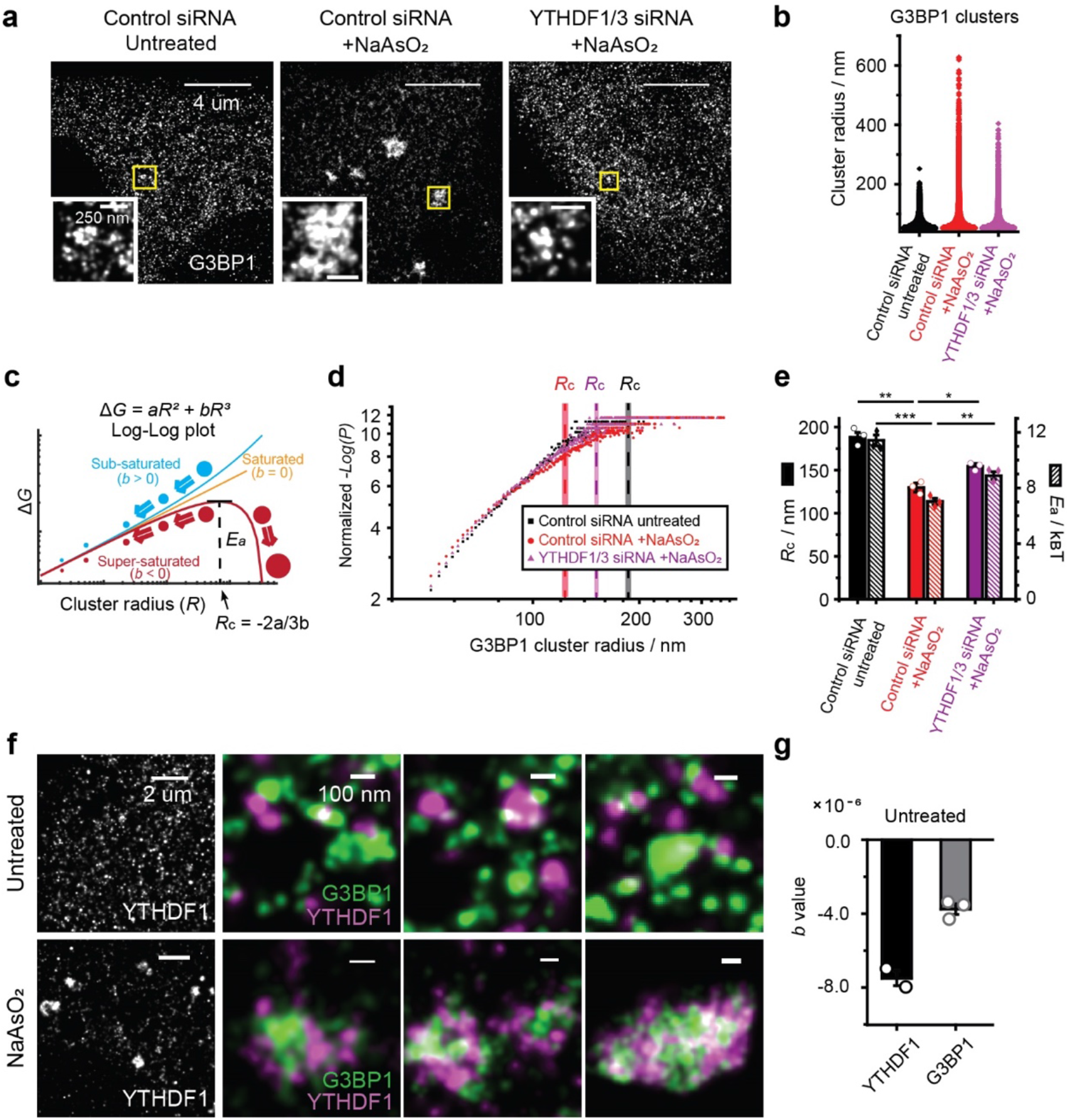
YTHDF proteins enhance phase-separation of G3BP1 by reducing the critical size and activation energy barrier for condensate formation. **a**, STORM imaging of G3BP1 protein in U-2 OS cells treated with control siRNA or YTHDF1/3 siRNA. The left panel shows an image of an unstressed cell treated by control siRNA, the middle panel shows an image of a NaAsO_2_-treated cell treated with control siRNA, and the right panel shows the image of a NaAsO_2_-treated, YTHDF1/3 knockdown cell. Insets are zoom-in images of regions in yellow boxes. **b**, Distribution of cluster radius for G3BP1 proteins under the three different conditions described in (**a**). More than 160000 clusters from ~60-80 cells from three independent experiments were pooled and analyzed for each condition. Counts of clusters with a radius larger than 50 nm are displayed. **c**, Diagram of Gibbs free energy change (Δ*G*) for cluster formation as a function of the cluster radius (*R*) for sub-saturated (*b* > 0), saturated (*b* = 0), and super-saturated (*b* < 0) states in the classical nucleation theory. Δ*G* contains two terms - a surface energy term and a bulk energy term: Δ*G* = *aR*^2^ + *bR*^3^. When molecules are in the super-saturated state, a critical radius (*R*_c_ = −2*a* / 3*b*) exists, beyond which the cluster continues to grow in size irreversibly. *E*_a_, the value of Δ*G* at *R*_c_, represents the activation energy barrier for super-critical cluster (condensate) formation. **d**, Log-log plots of normalized −Log(*P*) vs. *R* calculated from the size distribution of G3BP1 clusters show that G3BP1 is in a super-saturated state in unstressed cells (blacked), NaAsO_2_-treated cells (red) and NaAsO_2_-treated, YTHDF1/3 knockdown cells (magenta). *P* is the probability density of the clusters with radius *R*. In a steady-state system, *P* of sub-critical clusters (*R* < *R*_*c*_) follow Boltzmann distribution: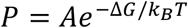. Thus, *ΔG* (*in k*_B_*T*) = −Log(*P*) − *c*, where *c* = −Log*A*. By fitting the data of −Log(*P*) versus *R* to the equation of −*Log*(*P*) = *aR*^2^ + *bR*^3^ + *c*, we obtained the values of *a*, *b*, from which we calculated the corresponding *R*_c_ and *E*_a_. The normalized values of −Log(*P*), which equal to −Log(*P*) − *c* and are equivalent to Δ*G* for sub-critical clusters, are plotted here. Data for clusters with radii between 50 nm and 350 nm are shown. The plot from one of the three independent experiments are shown here, and all three independent experiments show similar plots. Mean values of calculated critical radius *R*_c_ are shown as dashed lines with the shaded areas representing SEM for the 3 independent experiments. **e**, The critical radius (*R*_c_) and energy barrier (*E*_a_) for super-critical cluster (condensate) formation of G3BP1 derived from the measured cluster size distributions for the three conditions. Mean ± SEM are shown (n = 3 independent experiments for each condition). *: p < 0.05, **: p < 0.01, ***: p < 0.001, performed by unpaired two-tailed Student’s T-Test. **f**, Single-color STORM imaging of YTHDF1and two-color STORM imaging of YTHDF1 and G3BP1 in unstressed (upper panel) and NaAsO_2_-treated cells (lower panels). **g**, The *b* values in Δ*G* for YTHDF1 (n = 2 independent experiments) and G3BP1 (n = 3 independent experiments) clusters in unstressed U-2 OS cells.

We then investigated the effect of YTHDF proteins on the formation of G3BP1 clusters in the framework of the classical nucleation theory for first-order phase transitions^41,42^. In this model, the Gibbs free energy change (Δ*G*) for the formation of clusters with a specific radius (*R*) contains two terms - a surface energy term and a bulk energy term: Δ*G* = *aR*^2^ + *bR*^3^ (Fig. 6c). Three states can be discriminated using this model (Fig. 6c): sub-saturated state (*b* > 0), saturated state (*b* = 0), and super-saturated state (*b* < 0). In the super-saturated state, clusters that fluctuate to a critical size *R*_c_ (i.e., clusters that reached the activation energy barrier height *E*_a_) will continue to grow irreversibly and form super-critical clusters (Fig. 6c). Based on the distribution of G3BP1 cluster sizes measured by super-resolution imaging (Fig. 6d), we obtained the Δ*G* values for different cluster sizes (*R*), which allowed us to derive the values of *a* and *b*, as well as the values of *E*_a_, and *R*_c_ for G3BP1 clusters. Interestingly, G3BP1 appeared to be in a super-saturated state with a negative value of *b* even in unstressed cells (Fig. 6d), and NaAsO_2_-induced stress pushed G3BP1 into a deeper super-saturated state with a more negative *b* value (i.e. smaller *E*_a_ and *R*_c_, Fig. 6e). Notably, knockdown of YTHDF1/3 increased *R*_c_ and *E*_a_ (Fig. 6e), which in turn resulted in a decrease in the cluster sizes of G3BP1.

### YTHDF1 forms clusters connecting G3BP1 core clusters in SGs

Next, we performed STORM imaging on the endogenous YTHDF1 protein in U-2 OS cells. We observed that YTHDF1 also formed clusters in the unstressed condition and two-color STORM imaging of YTHDF1 and G3BP1 showed that the YTHDF1 and G3BP1 clusters did not substantially colocalize in unstressed cells (Fig. 6f, upper panels). Notably, analysis of the size distribution of the YTHDF1 clusters showed that YTHDF1 protein was also in a super-saturated state in unstressed cells, with a negative *b* value of even a greater magnitude than that of G3BP1 in unstressed cells (Fig. 6g). Upon NaAsO_2_ treatment, the sizes of YTHDF1 clusters increased significantly and many YTHDF1 clusters coalesced with G3BP1 clusters (Fig. 6f, lower panels). Interestingly, YTHDF and G3BP1 proteins did not mix completely in SGs; instead, the YTHDF clusters often resided on the periphery of individual G3BP1 clusters and at the junction connecting neighboring G3BP1 clusters.

## Discussion

In this study, we found that m^6^A-modified mRNA enriches in SGs, and the m^6^A-binding YTHDF proteins play a critical role in SG formation by reducing the critical radius and energy barrier for the phase transition of SG core proteins.

We showed that endogenous YTHDF1 and YTHDF3 were exclusively enriched in SGs but not P-bodies, while endogenous YTHDF2 was enriched in both SGs and P-bodies. Knockdown of YTHDF1/3 strongly inhibited SG formation and localization of mRNAs to SGs. Our results are in stark contrast with a recent report showing that SG formation is not affected by YTHDF1 or YTHDF3 knockdown^43^. However, in this previous work SGs are imaged through the expression of GFP-labeled G3BP1 in cells, and overexpression of this SG core protein likely has masked the effect of YTHDF proteins in SG formation.

Notably, we found both the N-terminal IDR and C-terminal m^6^A-binding YTH domains to be critical for promoting SG formation. The N-terminal IDR of the YTHDF proteins are Tyr(Y)-rich, and Arg(R)-deficient. These Tyr(Y) residues could interact with the Arg(R) residues in the YTH domain through π-cation interaction^44^ to promote clustering of YTHDF proteins. The Tyr(Y) residues in the N-terminal IDR of YTHDF proteins may also mediate interactions with other SG-components that are Arg(R)-rich such as eIF3A or other RNA-binding proteins^17^. Interestingly, the center PLDs of the YTHDF proteins are Pro(P)/Gln(Q)-rich, but Gly(G)-poor, which makes this region relatively rigid^44^. This rigid linker between the N-terminal IDR and C-terminal YTH domain could serve to prevent intramolecular π-cation interactions, and thus promote intermolecular interactions among YTHDF proteins, as well as between YTHDF and other SG proteins, thereby promoting the formation of protein condensates. We also found the m^6^A-binding activity of YTHDF proteins to be important for SG formation.

Furthermore, using super-resolution imaging, we found that both YTHDF1 and G3BP1 are in the super-saturated state in cells even in unstressed conditions. The super-saturated state of G3BP1 and YTHDF1 makes them ready to form super-critical clusters, which could ensure a sensitive response to environmental changes. However, according to the Szilard model of non-equilibrium steady-state super-saturation^42^, super-critical clusters need to be constantly removed to maintain a steady state in unstressed conditions, which could be mediated through autophagy^45^ or protein-RNA disaggregases^12^. Notably, YTHDF1/3 reduced the activation energy barrier for super-critical cluster (condensate) formation of the G3BP1, thereby promoting SG assembly in cells. Interestingly, YTHDF1 clusters tended to reside on the periphery of G3BP1 clusters and at the junctions connecting G3BP1 clusters, which can promote SG formation by connecting small G3BP1 core clusters into larger granules. It has been proposed that SGs adopt a heterogeneous structure formed by initial nucleation of the G3BP-rich cores followed by juxtaposition of the nucleated cores, potentially through a more dynamic shell^12,46^, but the identity of the SG shell protein(s) remains elusive. Our results suggest that YTHDF proteins function as SG-shell proteins that promote SG formation by bringing together multiple SG-core clusters to form large granules. Overall, our results provide new insights into the function of RNA modifications and RNA modification recognition proteins in regulating phase separation and membraneless compartment formation in cells.

## Acknowledgments

We thank members of Zhuang Lab for the kind help, especially Ruobo Zhou and Boran Han for help with two-color STORM setup and data analysis, Guiping Wang and Monica Thanawala for help with data analysis. We thank Dr. Ke Xu (University of California, Berkeley) for help with the script for two-color STORM data analysis. We thank Dr. Paul Anderson and Dr. Nancy Kedersha (Harvard Medical School) for helpful discussions, and Dr. Yang Shi (Harvard Medical School) for providing U-2 OS-METTL3-KO cell line. This work is in part supported by NIH. XZ is an HHMI investigator.

## Author contributions

YF and XZ designed the experiments. YF performed experiments and analyzed data. YF and XZ wrote the paper.

