## Supplementary Materials for "m^6^A-binding YTHDF proteins promote stress granule formation by modulating phase separation of stress granule proteins"

### Supplementary Figures

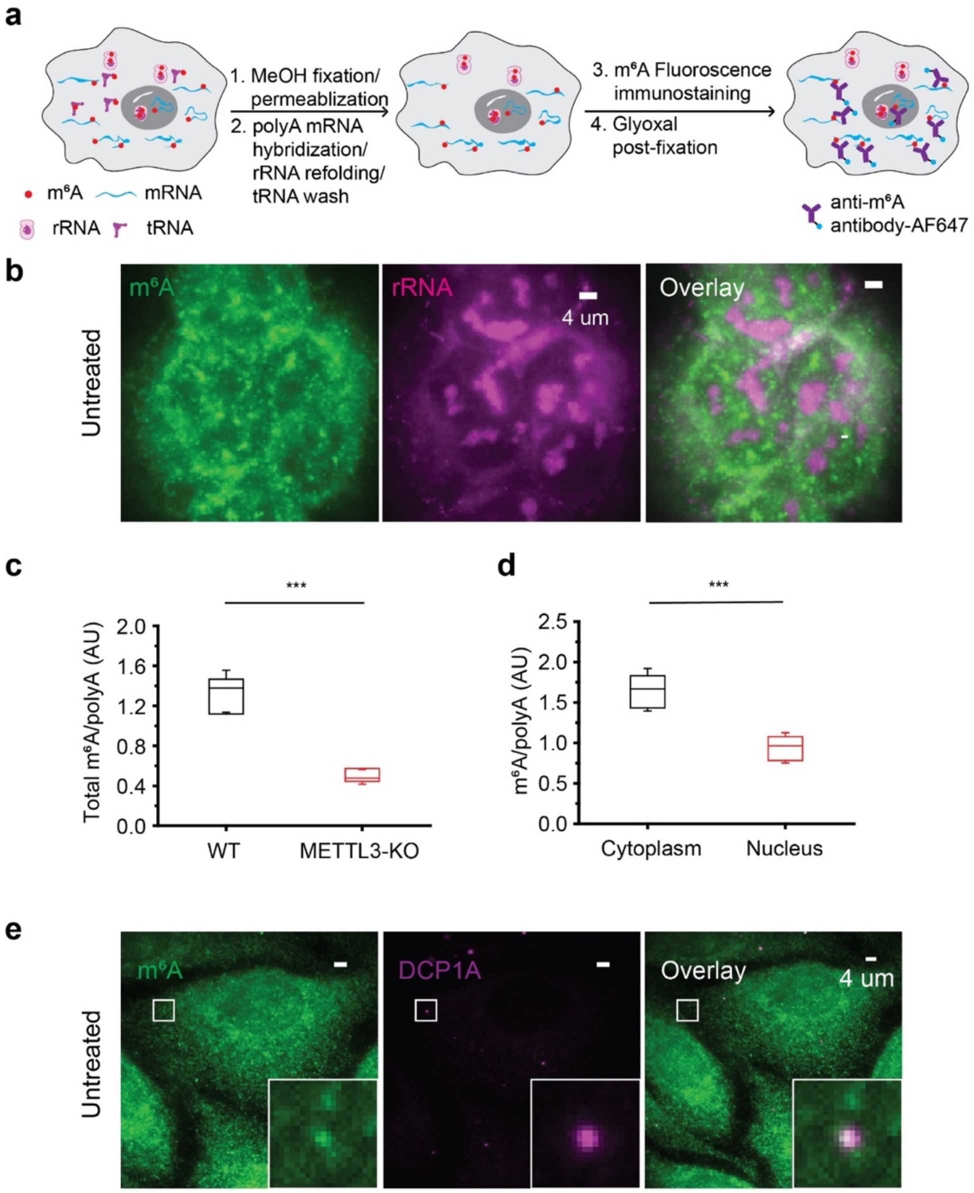


**Supplementary Fig. 1 | Immunofluorescence staining of m$^6$A in U-2 OS cells. a,** Schematic procedure for immunofluorescence staining of mRNA m$^6$A in fix cell. Methanol is used for cell fixation and permeabilization. Methanol fixation retains large RNAs, including mRNAs and rRNAs, but not small RNAs like tRNAs and snRNAs, which require strong covalent (such as aldehyde-based) fixation to be preserved in cells. Following methanol fixation, the polyA tails of the cellular RNAs are hybridized with poly-dT FISH probes, during which step the rRNAs are refolded, preventing the antibody binding to rRNA m$^6$A. Cells are then stained using AF647 labeled anti-m$^6$A polyclonal antibody and post-fixed with glyoxal solution. **b,** Two-color imaging of the $m^6A$ signal, detected by the labeling protocol described in (a), and the rRNA signal (detected by immunofluorescence staining with anti-rRNA antibody) shows that the $m^6A$ signal does not colocalize with the rRNA signal in the nucleolus in the above staining procedure. **c,** Normalized immunofluorescence signals of mRNA $m^6A$ in wildtype (WT) and METTL3-KO U-2 OS cells show around ~60% reduction of the $m^6A$ signal. The $m^6A$ signal is normalized against the polyA signal. n = 15 cells, from three independent experiments under each condition. **d,** The normalized mRNA $m^6A$ signal is higher in the cytoplasm than in the nucleus, consistent with a role of $m^6A$ in promoting mRNA export from cell nucleus, n = 14 cells, from three independent experiments. In **c** and **d**, box plots show the median, 25%-75% quartiles, and standard deviation. *** $P < 0.001$ performed by unpaired Mann-Whitney U Test. **e,** In unstressed U-2 OS cells, immunofluorescence staining of mRNA $m^6A$ shows a diffusive pattern of $m^6A$ signal in the cytoplasm and enrichment of $m^6A$ signal in P-bodies. P-body marker DCP1A is detected using immunofluorescence.

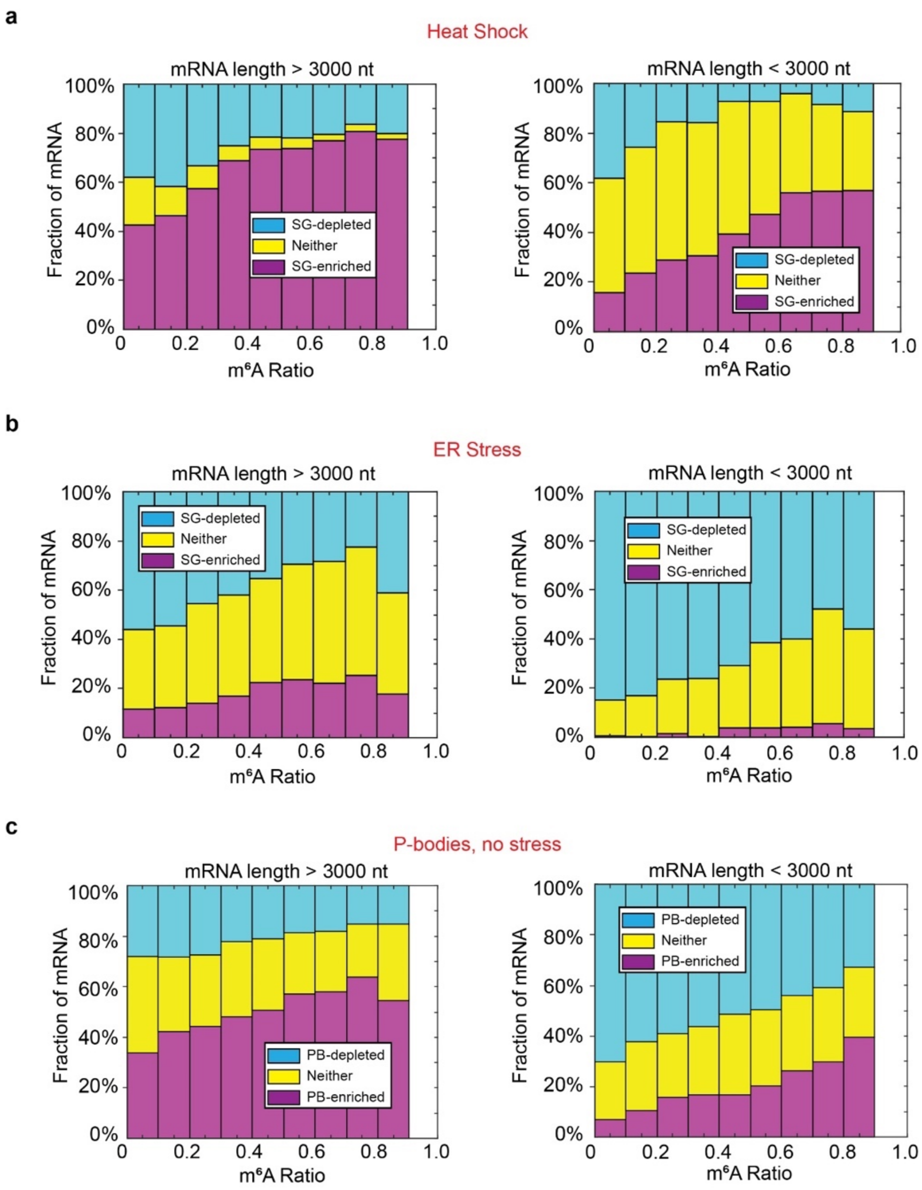


**Supplementary Fig. 2 | $m^6A$-modified RNAs are enriched in different types of SGs and in P-bodies. a, b, c,** Transcriptome-wide analysis of the relationship between the $m^6A$ ratio and SG (or P-body) enrichments in heat-shock induced SGs (a), in ER-stress induced SGs (b), and in P-bodies in unstressed cells (c). The fraction of mRNAs shown in magenta, cyan, and yellow bars are as defined as in Fig. 1b, c, but for heat-shock induced SGs, ER-stress induced SGs, and P-bodies, instead of oxidative-stress induced SGs. The $m^6A$ ratio is as described in Fig. 1b, c. mRNA species were separated into two categories for analysis: > 3000 nt (left panels) and < 3000 nt (right panels). n = 9049 genes in total.

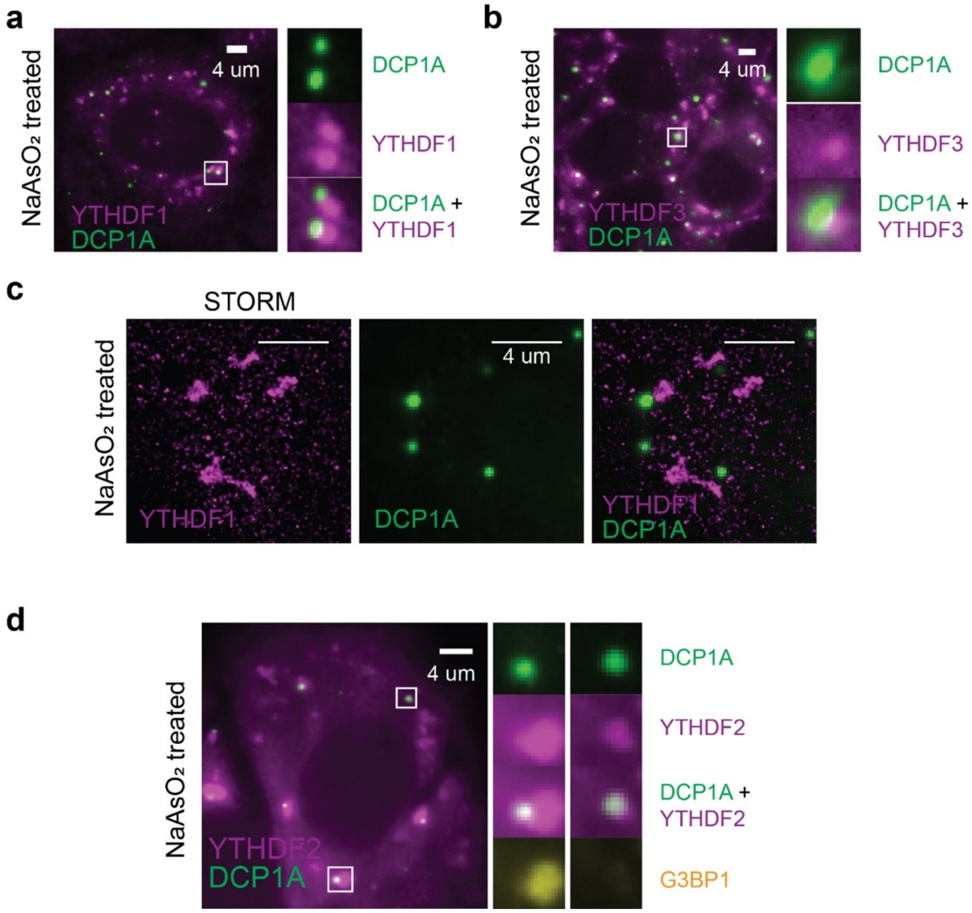


**Supplementary Fig. 3 | YTHDF2 but not YTHDF1 or YTHDF3 colocalizes P-bodies.**

**a, b,** Conventional immunofluorescence images of endogenous YTHDF1/3 and the P-body marker DCP1A in cells treated with 0.5 mM $NaAsO_2$ for 30 min to induce oxidative stress showing that YTHDF1 and YTHDF3 often lie adjacent to P-bodies. **c,** STORM imaging of endogenous YTHDF1, in combination with conventional imaging of DCP1A, confirms the absence of YTHDF1 from DCP1A marked P-bodies under oxidative stress. **d,** Conventional immunofluorescence images of endogenous YTHDF2 and the P-body marker DCP1A, as well as the SG marker G3BP1, in cells treated with 0.5 mM $NaAsO_2$ for 30 min. YTHDF2 colocalizes with P-body under oxidative stress, and G3BP1 is absent in the regions where YTHDF2 and DCP1A colocalize.

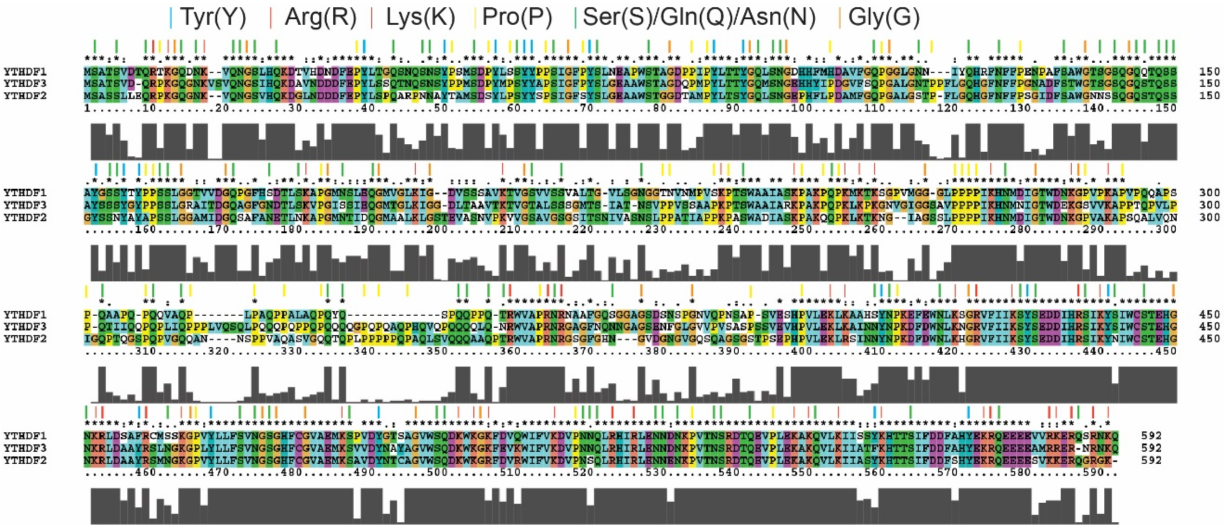


**Supplementary Fig. 4 | Sequence alignment of human YTHDF1, YTHDF2, and YTHDF3 proteins.** Human YTHDF1-3 protein sequences were aligned using ClusterX. Conserved amino acids that are important for prion-like domain (PLD) formation or mediate interactions important for phase separation are marked with colored sticks.

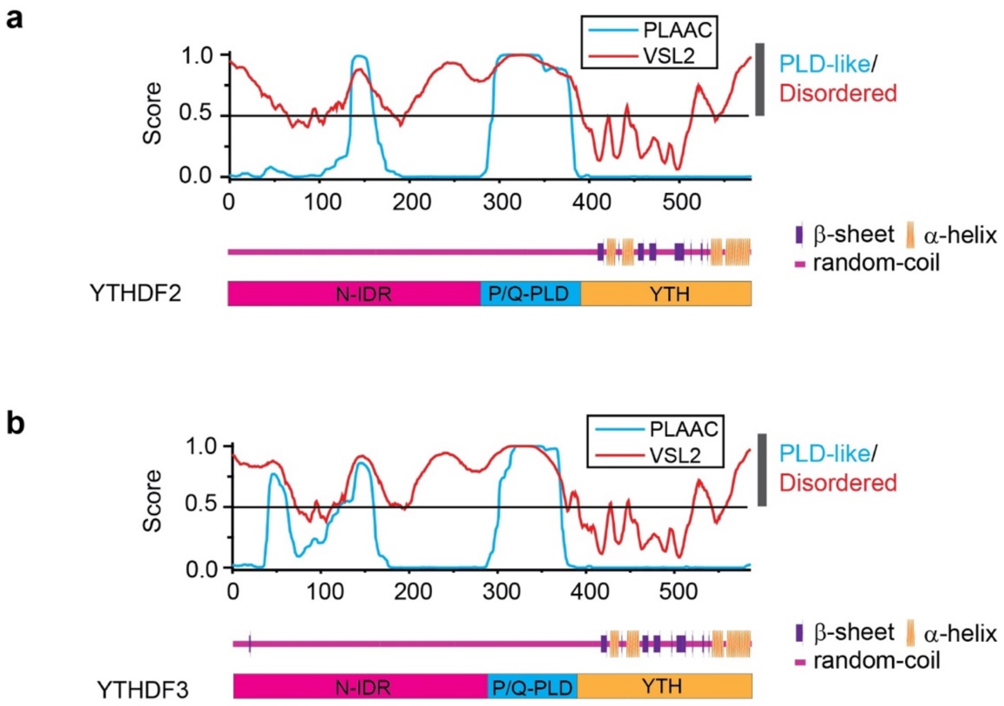


**Supplementary Fig. 5 | Amino acids composition and predictions of IDRs, PLDs, and secondary structures in YTHDF2 and YTHDF3 proteins. a, b,** The likelihood scores for being disordered or PLD-like, and the secondary structures, are predicted as described in Fig. 4a, but for YTHDF2 (a) and YTHDF3 (b) instead of YTHDF1.

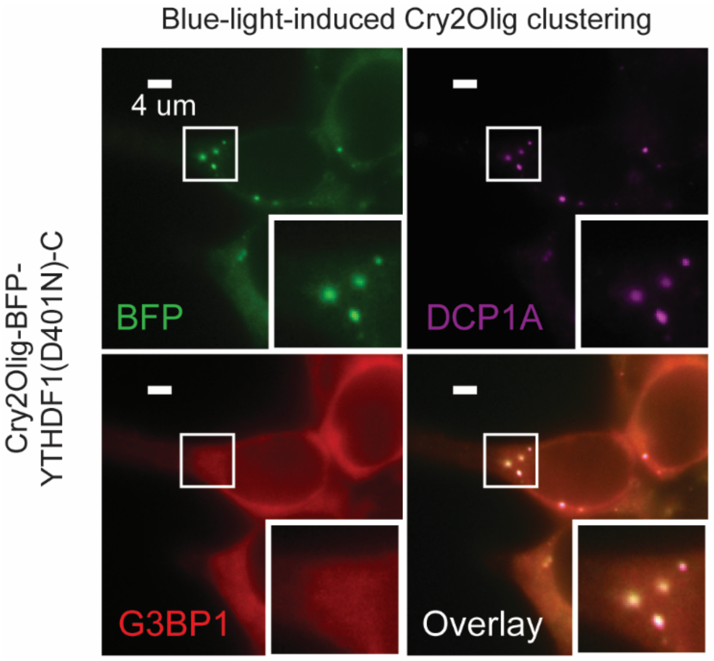


**Supplementary Fig. 6 | Light-induced oligomerization of proteins containing m$^6$A-binding YTH domain does not induce SG formation in the absence of stress.** Blue-light-induced oligomerization of Cry2Olig-BFP-YTHDF1(D401N)-C causes its colocalization with P-bodies, but does not induce the formation of SGs, in unstressed cells.

### Methods

#### Cell lines

U-2 OS cells (ATCC, HTB-96™), U-2 OS-METTL3 knockout cells (Yang Shi Lab) were grown at 37 °C and 5% $CO_2$ in EMEM medium supplemented with 100 U/mL streptomycin, 100 ug/mL penicillin and 10% fetal bovine serum. Cell medium was changed 24 hr before $NaAsO_2$ treatment. All imaging experiments were performed on cells plated on 20 mm coverslips in 12-well tissue culture plates, with density ranging from $8 \times 10^4$ cells/well to $1.6 \times 10^4$ cells/well depending on different treatment procedures.

#### Antibodies

All fluorescent-dye-conjugated primary antibodies were labeled using Succinimidyl (NHS) esters following manufacture's protocols and stored in PBS with 40% glycerol at -80 °C. For two-color STORM imaging, primary antibodies were labeled using CF®680 Succinimidyl esters (Biotium, 92139) or AlexaFluor 647-NHS esters (ThermoFisher, A37573). Mouse monoclonal anti-Digoxigenin antibody (Abcam, Ab420) was labeled using CF®750 Dye Succinimidyl Ester (Biotium, 92142). The following antibodies were used in this study: Rabbit polyclonal anti-$m^6A$ antibody (Abcam, ab151230); Rabbit polyclonal anti-$m^6A$ antibody, Alexa Fluor® 647-conjugated (labeled using ab151230, 1:4 labeling ratio); Rabbit monoclonal anti-G3BP antibody, Alexa Fluor® 647 conjugated (Abcam, ab215944); Rabbit monoclonal anti-G3BP antibody, Alexa Fluor® 555 conjugated (Abcam, ab217729); Mouse monoclonal anti-G3BP Antibody (Abcam, ab56574); Mouse monoclonal anti-G3BP Antibody, CF®680 conjugated (labeled using ab56574, 1:5 labeling ratio); Rabbit monoclonal anti-DCP1A antibody, Alexa Fluor® 488 conjugated (Abcam, ab208275); Mouse monoclonal rRNA Antibody (Y10b), Alexa Fluor® 488 conjugated (Novus Biologicals, NB100-662AF488); Mouse monoclonal anti-Digoxigenin antibody CF750 conjugated (labeled using Abcam, Ab420, 1:5 labeling ratio); Rabbit Polyclonal anti-YTHDF1 Antibody (Proteintech, 17479-1-AP); Rabbit polyclonal anti-YTHDF1 Antibody CF®680 conjugated (labeled using 17479-1-AP, 1:5 labeling ratio); Rabbit monoclonal anti-YTHDF2 antibody (Abcam, ab245129); Rabbit monoclonal Anti-YTHDF2 antibody CF®680 conjugated (labeled using ab245129, 1:6 labeling ratio); Rabbit polyclonal anti-YTHDF3 Antibody (Proteintech, 25537-1-AP). The YTHDF3 antibody is detected by Alexa Fluor® 647 conjugated Goat anti-Rabbit IgG (H+L) secondary antibodies (ThermoFisher, A-21245).

#### Cloning and LentiVirus production

YTHDF protein constructs were cloned into $pSNAP_f$ vectors (New England BioLabs, N9183S) between AscI and EcoRI restriction enzyme cutting sites using standard cloning method. SNAP-tag was at the C-terminus of the construct. Plasmid constructs name and sequences are as following: pSNAPf-YTHDF1 (CCDS ID: CCDS13511.1), pSNAPf-YTHDF2 (CCDS41296.1), pSNAPf-YTHDF3 (CCDS75747.1), pSNAPf-YTHDF1-N (1-359 from CCDS13511.1), pSNAPf-YTHDF2-N (1-381 from CCDS41296.1), pSNAPf-YTHDF3-N (1-385 from CCDS75747.1), pSNAPf-YTHDF1-C1 (262-559 from CCDS13511.1), pSNAPf-YTHDF1-C2 (360-559 from CCDS13511.1).

For the Cry2Olig-BFP plasmid, Cry2Olig sequence from plasmid CRY2olig-mCherry (Addgene, 60032) was cloned into pFUGW together with a TagBFP sequence using Gibson assembly. A YTHDF1 fragment (264-559 from CCDS13511.1) with D401N mutation was then cloned into the Cry2Olig-BFP plasmid to yield Cry2Olig-BFP-YTHDF1(D401N)-C. The plasmids were then packed into lentivirus using the Lenti-X™ Packaging Single Shots (VSV-G) transfection kit (Clontech, 631276) in a Lenti293T cell line.

**Knockdown and overexpression of YTHDF proteins**

Transfection of siRNA, and co-transfection of siRNA and plasmids containing YTHDF proteins, were performed using Lipofectamine 3000 (ThermoFisher, L3000008) according to the manufacturer's protocol. The siRNAs we used here for YTHDF1/3 target 5'- or 3'-UTR regions of mRNAs and do not target the mRNA transcripts from transfected plasmids. siRNA used here are: YTHDF1 siRNA (Qiagen, SI00764715), YTHDF2 siRNA (Qiagen, SI04174534), YTHDF3 siRNA (Qiagen, SI00764778), AllStars Negative Control siRNA (Qiagen, SI03650318). SNAP-tagged YTHDF proteins were labeled using SNAP-Cell® TMR-Star (New England BioLabs, S9105) after cell fixation and permeabilization according to the manufacturer's procedure, followed by immunofluorescence staining using anti-G3BP1 antibodies.

**Induction of oligomerization of Cry2Olig constructs in cells**

After cells were infected by Lenti-virus containing Cry2Olig-BFP-YTHDF1(D401N)-C construct for 48 hours, oligomerization of Cry2Olig in cells was induced by exposing cells to a Maestrogen 470 nm UltraBright LED Transilluminator for 2 min at room temperature. Cells were subsequently fixed to perform immunofluorescence staining.

**Simultaneous polyA FISH and m$^6$A immunofluorescence staining**

We used methanol (MeOH) fixation to retain large RNAs, including mRNAs and rRNAs, but not small RNAs like tRNA and snRNA, which require strong covalent (such as aldehyde-based) fixation to be preserved in cells[47]. 3D structures of ribosomes showed that m$^6$A in both 18S and 28S rRNA reside inside well-folded rRNA structures[48,49]. Although MeOH denatures rRNA and exposes the m$^6$A site in rRNA, we included a refolding step to refold rRNAs and prevent the binding of rRNA m$^6$A by the anti-m$^6$A antibodies. Meanwhile, although anti-m$^6$A antibody can also recognize m$^6$A in long non-coding RNA (lncRNA) and *N*$^6$-2-*O*-dimethyladenosine (m$^6$A$_m$) in the 5' of mRNA, the amount of m$^6$A in lncRNA and the amout of m$^6$A$_m$ are less than 5% of total m$^6$A in mRNA[29]. Thus, our m$^6$A-immunostaining protocol primarily detects mRNA m$^6$A in mammalian cells.

Cells were grown on 20 mm coverslips in 12-well tissue culture plates. The cells were washed once with 1 mL of PBS and fixed and permeabilized with 1 mL of MeOH at -20 °C for 8-10 min. After withdrawing MeOH, cells were dried completely under air for 5-10 min and equilibrated in 1 mL of Stellaris® RNA FISH Wash Buffer A (LGC Biosearch Technologies, SMF-WA1-60) containing 10% formamide for 5 min. RNA FISH hybridization solution was prepared using 1 uM of DIG-polydT-LNA probe (Sequence: /5DigN/T+TT+TT+TT+TT+TT+TT+TT form IDT, +T represents LNA form of T) in Stellaris® RNA FISH Hybridization Buffer (LGC Biosearch Technologies, SMF-HB1-10) containing 10% formamide. The coverslips were flipped to cover 30 uL of RNA FISH hybridization solution on a parafilm placed in a plate, and then incubated in a humidified 37 °C incubator overnight. The next day, coverslips were washed with 1 mL of Stellaris® RNA FISH Wash Buffer A twice for 30 min each at 37 °C. Coverslips were then washed and equilibrated three times with 1 mL of PBS at room temperature for 5 min each. Blocking solution was prepared as the following: 1× PBS, 2% UltrapureBSA (ThermoFisher, AM2616), 0.05% Triton-X100 (Sigma-Aldrich, T9284), 1:100 RNasin® plus (Promega, N2611) in RNase-free $H_2O$. The coverslip was blocked for 1 hr at room temperature with 15 uL of blocking solution on parafilm in a plate with wet Kimwipes on the side to prevent solution evaporation. Meanwhile, antibody solution was prepared by adding the following antibodies to 20 uL of block solution: CF750 conjugated anti-Digoxigenin antibody (1:50 dilution from 0.5 mg/mL stock), Alexa Fluor® 647 conjugated anti-$m^6A$ antibody (1:30 dilution from 0.4 mg/mL stock), Alexa Fluor® 555 conjugated anti-G3BP antibody (1:200 dilution from 0.5 mg/mL stock), Alexa Fluor® 488 conjugated anti-DCP1A antibody (1:200 dilution from 0.5 mg/mL stock) and/or Alexa Fluor® 488 conjugated anti-rRNA antibody (1:100 from 0.5 mg/mL stock). The coverslips were lifted from parafilm after the 1 hr blocking, and 20 uL of the antibody solution was added before the coverslips were put back to cover the solution. The incubation was performed overnight at 4 °C in the dark. The next day, coverslips were washed in a 12-well plate with 0.05% Triton-X100 in PBS for 4 times with 4 min each. Finally, coverslips were fixed with a 3% Glyoxal fixation solution[50] for 30 min at room temperature and washed for 3 times using PBS. The ~4 ml glyoxal fixation solution contained 2.835 ml $ddH_2O$, 0.789 ml ethanol (absolute, for analysis), 0.313 ml glyoxal (40% stock solution), 0.03 ml acetic acid, and adjust to pH 5 with 5M NaOH.

**Simultaneous mRNA smFISH and immunofluorescence staining**

For simultaneous mRNA smFISH and immunofluorescence staining, cells were first fixed with 4% Paraformaldehyde in PBS for 5 min, and then permeabilized with MeOH at -20 °C for 8-10 min, dried and equilibrated with Stellaris® RNA FISH Wash Buffer A containing 30% formamide. 0.5 uM of RNA smFISH probes with 30 nt hybridization region and a 20 nt readout sequence were used in the Stellaris® RNA FISH Hybridization Buffer with 30% formamide. After wash, coverslips were further hybridized to fluorophore-labeled readout probes (/5Alexa750N/ACACTACCACCATTTCCTAT or /5ATTO565N/ACCACAACCCATTCCTTTCA, IDT), complementary to the readout sequences on the smFISH probes, and washed twice with Stellaris® RNA FISH Wash Buffer A containing 10% formamide for 30 min at 37 °C. Coverslips were then washed and equilibrated three times with 1 mL of PBS at room temperature for 5 min each. Blocking solution was prepared as the following: 1× PBS, 2% UltrapureBSA (ThermoFisher, AM2616), 0.05% Triton-X100 (Sigma-Aldrich, T9284), 1:100 RNasin® plus (Promega, N2611) in RNase-free $H_2O$. Cells were blocked for 1 hr at room temperature with 15 uL of blocking solution on parafilm in a plate with wet Kimwipes on the side to prevent solution evaporation. Meanwhile, antibody solution was prepared by adding the following antibodies to 20 uL of block solution: Alexa Fluor® 555 conjugated anti-G3BP antibody (1:200 dilution from 0.5 mg/mL stock), and Alexa Fluor® 488 conjugated anti-DCP1A antibody (1:200 dilution from 0.5 mg/mL stock). The incubation was performed overnight at 4 °C in the dark. The next day, coverslips were washed with 0.05% Triton-X100 in PBS for 4 times with 10-15 min each. Finally, coverslips were fixed with 4% Paraformaldehyde in PBS for 30 min at room temperature and washed for 3 times using PBS.

**Two-color immunofluorescence staining for STORM imaging**

Cells on 20 mm coverslip were fixed with 3% Glyoxal fixation solution[50] for 15 min at 4 °C and 15 min at room temperature. Cells were then quenched with 50 mM of $NH_4Cl$ for 20 min at room temperature, and permeabilized with 0.5% (v/v) Triton X-100 in PBS for 4 min. Cells were then blocked with 3% BSA in PBS, and inverted on a parafilm with solutions containing with Alexa Fluor® 647 conjugated anti-G3BP1 antibody (1:30 dilution from 0.5 mg/mL solution) and CF680-conjugated anti-YTHDF1 antibody (1:30 dilution from 0.5 mg/mL solution) in 30 uL of 3% BSA in PBS at 4 °C overnight. The cells were then washed 4 times with 0.05% Triton-X100 in PBS for 10 min each. Cells were post-fixed with 4% Parafomaldehyde + 0.1% Glutaraldehyde in PBS for 30 min at room temperature, washed 3 times with PBS, and stored in PBS at 4 °C before STORM analysis.

**STORM imaging procedure**

For STORM imaging, we used a custom-built microscope with an Olympus IX-71 inverted microscope body, a UPlanSApo 100× N.A. 1.40, oil-immersion objective (Olympus), and an active auto-focusing system consisting of an infrared 830 nm laser (LPS-830-FC, Thorlabs) and a quadrant photodiode as described previously[51]. A 640-nm laser (Coherent) was used to excite and image Alexa Fluor® 647 (AF647) and CF680 on the antibodies. And a 405 nm laser (CUBE 405-50C, Coherent) was used to activate the fluorophores. The lasers were directed to the sample using a dichroic mirror (ZET405/488/561/640mv2, chroma) on the excitation path. For two-color imaging of AF647 and CF680, on the emission path, we used a Dual-View setup (DV-CC, Dual-view, Photometrics) with a 685-nm long-pass dichroic mirror (Chroma, T685lpxr) to separate the emission wavelengths and projected the emission photons to two separated regions on EMCCD camera. The two Dual-View channels were aligned by taking calibration images of 100-nm Tetraspeck beads (Invitrogen) attached to a coverslip surface. Illumination was adjusted to near total internal reflection fluorescence configurations.

Cells on a 12 mm coverslip were assembled on a cover-slide and sealed in imaging buffer. The imaging buffer contains 200 mM cysteamine (Sigma), 5% glucose (Sigma), 0.8 mg/mL glucose oxidase (Sigma), and 40 µg/mL catalase (Roche Applied Science). During imaging, 640-nm laser (~2 kW/cm$^2$) was used to excite AF647 and CF680 to switch them into the dark state. A 405-nm laser was used to reactivate the fluorophores to the emitting state. The power of the 405-nm lasers (0 - 1 W/cm$^2$) was adjusted during image acquisition so that at any given instant, only a small, optically resolvable fraction of the fluorophores in the sample was in the emitting state.

**Analysis of correlation between SG enrichment and m$^6$A ratio**

For individual mRNA species, m$^6$A ratios only change slightly between cell types (most changes are less than 20%)[29], and $NaAsO_2$ treatment does not affect the methylation of mRNA that are methylated in CDS or 3'-UTR in U-2 OS cells[43]. We also found that $NaAsO_2$-treatment does not change the overall m$^6$A level in mRNA as measured by UHPLC-QQQ-MS/MS. Thus, we performed the correlation analysis using the available mRNA m$^6$A ratio data from two human cell lines[29], and combined them with the available datasets for mRNA enrichment in $NaAsO_2$-induced SGs[28], mRNA enrichments in other types of SGs[32] and mRNA enrichment in P-bodies in U-2 OS cells[33]. All mRNA with available m$^6$A ratios and with FPKM (Fragments Per Kilobase of transcript per Million mapped reads) > 50 from U-2 OS mRNA sequencing data were analyzed here. To mitigate the effect of mRNA length, which can influence SG enrichment, we separated mRNAs into two categories: long mRNA (longer than 3000 nucleotides (nt) in total mRNA length), and short mRNA (shorter than 3000 nt). We then quantified the fractions of mRNA that are enriched in (> 2-fold enrichment in SGs or P-bodies), depleted from (< 0.5-fold enrichment), or neither enriched in nor depleted from (between 0.5- and 2-fold or statistically insignificant) SGs (or P-bodies) within different ranges of m$^6$A ratios.

**Quantification of m$^6$A and polyA immunofluorescence signal in SG**

Fluorescence images were analyzed using a custom MATLAB script. After background subtraction, particle analysis was performed on G3BP1 image to create the mask for the SG/non-SG region. Segmentation of individual cells and nucleus was then performed manually based on the polyA RNA staining signals. After subtracting the background, images of polyA and m$^6$A were quantified using the masks for SG/non-SG regions in the cytoplasm region of individual cells. Enrichment ratios of polyA and m$^6$A signals in SG were determined based on the average intensities of polyA and m$^6$A in the SG and non-SG regions in the cytoplasm.

**Data analysis for STORM imaging**

For single-color STORM, STORM movie was analyzed using a custom Insight3 software as previously described[52]. Briefly, fluorescence peaks of individual molecules were identified and fit to a 2D Gaussian to determine each peak's centroid position (x, y) and intensity. Sample drift during acquisition was subtracted. The resulting localizations were collected in a single molecule list file for further analysis.

For two-color STORM of AF647 and CF680, STORM movies were collected in two separate areas of the camera (256 × 256 pixels each) corresponding to two-color channels (>685 nm and <685 nm in wavelength), and analyzed using the above-described procedure to give a list of single molecule localizations for each channel. AF647 signals appear in both channels, and CF680 signals only appear in the > 685 nm channel. The signals from > 685 nm channel were assigned to AF647 or CF680 using the following procedure: The localization positions from two channels were first aligned based on the Tetraspeck bead signals using MATLab's cp2tform function with the 'projection' parameter. We then identified corresponding localizations from both channels within 1 pixel in the middle 5000 frames, and use them to further align the entire molecule list using MATLab's cp2tform function using 'polynomial' parameter. After the alignment, for each localization in > 685 nm channel, we search whether there is a corresponding localization in the <685 nm channel within 1 pixel, if so the localization is assigned to be correspondent to AF647. Otherwise, it is assigned to CF680. To further correct for any mis-assignment of CF680 molecules, which happens when a single-molecule signal on the <685 nm channel was not identified correctly mostly due to the overlap with an adjacent single-molecule signal, we also read the intensity of the corresponding pixels in the original movie and calculate the intensity ratios from two channels. Localizations with $(I_1-I_2)/(I_1+I_2) < 0.5$ were discarded, and only those with this value > 0.5 are kept as CF680 signals ($I_1$: intensity in > 685 nm channel, $I_2$: intensity in < 685 nm channel). Localizations from < 685 nm channel were automatically assigned to AF647 and analyzed together with the localizations that were assigned to AF647 from > 685 nm channel. Any misalignment can be identified from pattern mismatch of AF647 in these two categories of molecules.

Imaging resolutions are 22 nm for AF647 and 26 nm for CF680, as determined by the full-width-at-half-maximum of the localization distributions from all clusters in unstressed cells. Crosstalk ratios between two channels were determined to be less than 0.2% from AF647 to CF680, and 0.002% (20 ppm) from CF680 to AF647, which were determined by imaging samples labeled with AF647- or CF680- labeled antibodies separately.

**Cluster analysis from single molecule localization data**

A custom MATLAB script was developed to analyze the area of individual clusters. The localizations calculated from STORM movie were rendered into 2D-images by calculating the histogram with a 2D-bin of 15 nm by 15 nm. Particle analysis was performed to determine the area of individual clusters using a MATLAB thresholding function 'graythresh'[53], followed by functions of 'imbinarize', 'bwlabel', and 'regionprops'. Areas of all clusters were calculated. Typically 3000 clusters can be identified from each cell.

**Classical nucleation model**

Gibbs free energy change for cluster formation of clusters at a certain size (diamaeter $R$) is described by the equation: $\Delta G = 4\pi R^2\gamma - \frac{4}{3}\pi R^3\rho\Delta\mu$. In this equation, γ is the surface tension of clusters, ρ is the density of molecules inside clusters, $\Delta\mu = k_B T Log(\frac{c_{sol}}{c_{sat}})$ is the chemical-potential difference between molecules in the solution phase and in the cluster phase, in which $c_{sol}$ is the solution concentration of molecules, and $c_{sat}$ is the saturation concentration of molecules [42,54-56]. We abbreviate the equation as $\Delta G = aR^2 + bR^3$. In a steady-state system, the distribution probabilities of sub-critical clusters follow Boltzmann distribution: $P = Ae^{-\Delta G/k_B T}$, thus, $\Delta G = -Log(P) - c$ in the unit of $k_B T$, and $-Log(P) = aR^2 + bR^3 + c$, where $c = -LogA$[42]. When molecules are in the super-saturated state, a critical radius ($R_c = -\frac{2a}{3b}$) exists at $\frac{d\Delta G}{dR} = 0$. The activation free energy barrier $E_a$ was calculated from $\Delta G$ at $R_c$: $E_a = aR_c^2 + bR_c^3 + c$.

**Data fitting for classical nucleation theory model**

A group of 10-25 cells were pooled for analysis from a single experiment. Cluster volumes were calculated from the cluster area values by assuming sphere shape for individual clusters. Clusters volumes were binned with a bin size of $1 \times 10^5$ nm$^3$ from 523,099 nm$^3$ to 179,594,380 nm$^3$, corresponding to cluster radii from 50 nm to 350 nm. The frequency of cluster on each bin was calculated as *P*. *-LogP* (natural log of P) was plotted against the median radius *R* for the corresponding bin, which was calculated using the median volume of the corresponding bin. Parameters *a*, *b*, and *c* were determined by fitting a function $-LogP = aR^2 + bR^3 + c$ to the data points using MATLab's cftool function to 'poly3' with the parameter limit of $R^1$ set to 0. $R_c$ was calculated by $R_c = 2a/3b$, since only datapoints with a value less than $R_c$ can be used for the fitting, the fitting was iterated with subsets of data that eliminate datapoints with *R* larger than $0.8 \times R_c$, until *Rc* converges between two iterative rounds of fitting (i.e. the calculated *Rc* is larger than 1.25× the largest *R* in the data used for fitting). Values of *a*, *b*, *c*, $R_c$ and $Ea = aR_c^2 + bR_c^3 + c$ was reported. Their mean value and S.E.M. was calculated based on data fitting from three independent experiments.
